# Inferring the selective history of CNVs using a maximum likelihood model

**DOI:** 10.1101/2024.01.15.575676

**Authors:** Seyed Amir Malekpour, Ata Kalirad, Sina Majidian

## Abstract

Copy number variations (CNVs) – structural variations generated by deletion and/or duplication that result in change in DNA dosage – are prevalent in nature. CNVs can drastically affect the phenotype of an organism and have been shown to be both involved in genetic disorders and be used as raw material in adaptive evolution. Unlike single-nucleotide variations, the often large and varied effects of CNVs on phenotype hinders our ability to infer their selective advantage based on the population genetics data. Here, we present a likelihood-based approach, dubbed PoMoCNV, that estimates the evolutionary parameters of CNVs based on population genetics data. As a case study, we analyze the genomics data of 40 strains of *Caenorhabditis elegans*, representing four different populations. We take advantage of the data on chromatin accessibility to interpret the evolutionary parameters of CNVs inferred by PoMoCNV. We further test the reliability of PoMoCNV by estimating the evolutionary parameters of CNVs for mutation-accumulation experiments in *C. elegans* with varying levels of genetic drift.

**Significance:** Inferring the evolutionary parameters of copy number variations (CNVs) based on population genetics data is crucial to understand their role in evolution. However, given the diversity in the size and effects of CNVs, such inference poses a challenge. We developed a likelihood-based approach called PoMoCNV to address this issue.

## Introduction

Structural variations, i.e., genomic changes spanning more than 1 kilo bases (kb), has been widely recognized as a major source of genetic variability within and between species (. Copy number variations (CNVs) are a subset of structural variations that are specifically generated by duplication and deletion (Malekpour et al., 2018). Given the considerable length of CNVs, their effects on the phenotype, compared to single-nucleotide variations (SNVs), could be substantial. In fact, the ever-growing body of literature on CNVs in humans has delivered a hefty catalog of such variations in human genome (Feuk et al., 2006; Zarrei et al., 2015) and the myriad of diseases linked to this group of structural changes (Conrad et al., 2010; Dumas and Sikela, 2009; Tang and Amon, 2013; Zhang et al., 2009). Aside from such detrimental effects, the co-option of gene duplicates have long been regarded as a short path to the emergence of novel functions (Conant and Wolfe, 2008; Ohno, 1970; Ponting, 2008). In that respect, CNVs has been shown to provide just such raw material for adaptive evolution in a diverse array of organisms. For example, the duplication of hexose transporter in *Saccharomyces cerevisiae* confers adaptive benefit (i.e. higher fitness) in stressful conditions (reviewed in (Kondrashov, 2012)), and in humans, copy number of the salivary amylase gene seems to have increased in population accustomed to high-starch diets (Perry et al., 2007). Even in plants, CNVs appear to provide crucial genetic raw material for selection during domestication (Lye and Purugganan, 2019). Furthermore, the dazzling array of mind-powered innovations since the start of Holocene, usually attributed to the expansion of neocortex in our species can be traced back to the repeated duplication of *SRGAP2* in the lineage leading to *Homo sapiens* (Dennis et al., 2012).

In spite of their prevalence in nature, understanding the evolutionary dynamics of CNVs has proved a Herculean task. Whereas an SNV can be linked to a given biological functions, a CNV, given its length, would more likely be pleiotropic. Consequently, the available data on SNVs can be used to draw evolutionary conclusions, e.g., to understand compensatory evolution by linking SNVs to structural changes in proteins across phyla (Ivankov et al., 2014; Kondrashov et al., 2002), whereas making similar evolutionary inferences with respect to CNVs is not straightforward.

One widely-used approach to understand the fitness effects of SNVs has been to use their current frequencies in a population or populations, combined with *a priori* information on their demographic histories, to infer the patterns of selection in the past. This approach has been applied to the polymorphism data in humans to infer the evolutionary consequences of amino acid substitutions (Boyko et al., 2008; et al., 2005). De Maio *et al*. (De Maio et al., 2013) combined the available data on humans and their great ape relatives, chimpanzees and orangutans to infer their evolutionary history, i.e., fixation rate and mutation rate. In this contribution, we purpose a novel POlymorphism-aware phylogenetic MOdel (PoMo) for CNV datasets, dubbed PoMoCNV, which infers fitness parameters and the rate of CNV evolution based on the genomic data.

As a case study, we applied PoMoCNV to understand the evolution of CNVs in *Caenorhabditis elegans*. The sheer number of duplicated genes in *C. elegans* (*⇠* 8971) dwarfs those of *Saccharomyces cerevisiae* (*⇠* 1858) or *Drosophila melanogaster* (*⇠* 5536) (Rubin et al., 2000), with an estimated rate of duplication of *⇠* 0.02 per gene per million years, compared to *⇠* 0.002 in *D. melanogaster* (Lynch and Conery, 2000). Given such a high rate of gene duplication, *C. elegans* has been used to study the evolutionary dynamics of CNVs during laboratory experimental evolution. For example, (Konrad et al., 2018) took advantage of the short generation time of *C. elegans* – roughly 3 days (Wood, 1988) – and followed the emergence and loss of duplicates in a mutation accumulation (MA) experiment for *⇠* 400 generations. To vary the relative strength of selection to genetic drift, Konrad *et al*. used three different bottleneck sizes, *N* = 1, *N* = 10, and *N* = 100, to propagate different lines in the MA experiment. The dataset generated on the dynamics of the CNVs during this MA experiment, provides a perfect testing bed for *PoMoCNV*, since no speculation concerning the demographic history is required vis-à-vis phylogenetic inference and the strength of selection in each lineage is known *a priori*. In addition, the available in-depth knowledge concerning the genome regulation in *C. elegans*, specifically chromatin accessibility (Daugherty et al., 2017; Evans et al., 2016; Gerstein et al., 2010; Valouev et al., 2008), can be used to provide a more accurate inference of the evolutionary dynamics of CNVs.

## Results

PoMoCNV is employed to estimate the evolutionary parameters for the following *C. elegans* datasets: (i) CNVs in four *C. elegans* populations (Africa, Australia, France, and Hawaii), each consisting of 10 strains. Where each strain represents a specific *C. elegans* individual with unique genetic characteristics. For further details refer to (Lee et al., 2021), (ii) for CNVs in the aforementioned *C. elegans* populations, we utilized ATAC-seq (Assay for Transposase-Accessible Chromatin using sequencing) data (Jänes et al., 2018) to categorize chromatin into open or closed segments. Evolutionary rates are then estimated separately for the open and closed chromatin segments, (iii) CNVs in three MA experiments with bottleneck sizes *N* = 1, *N* = 10, or *N* = 100, in *C. elegans* populations (Konrad et al., 2018). These MA experiments allow for copy gain and loss events under varying selection intensities in *C. elegans*.

### CNV evolution rate estimation in *C. elegans*

The evolution rate of copy numbers is estimated using DNA sequencing data. We benefited from publicly available short read DNA sequencing data of *C. elegans* (Lee et al., 2021). The first step is to align reads to the reference genome (upper panel of Figure 1). We aligned the short read DNA sequences from four *C. elegans* populations (Africa, Australia, France, and Hawaii), each consisting of 10 strains, to the N2 reference genome (WS270). CNVs are then infered with CNVnator (Abyzov et al., 2011). After dividing the genome into non-overlapping segments with a length of 1kb, copy numbers, including homozygous and heterozygous deletions (zero and one copy), normal (2 copies), and duplications (more than 2 copies), are assigned to each segment using CNV calls from CNVnator.

**Figure 1.**
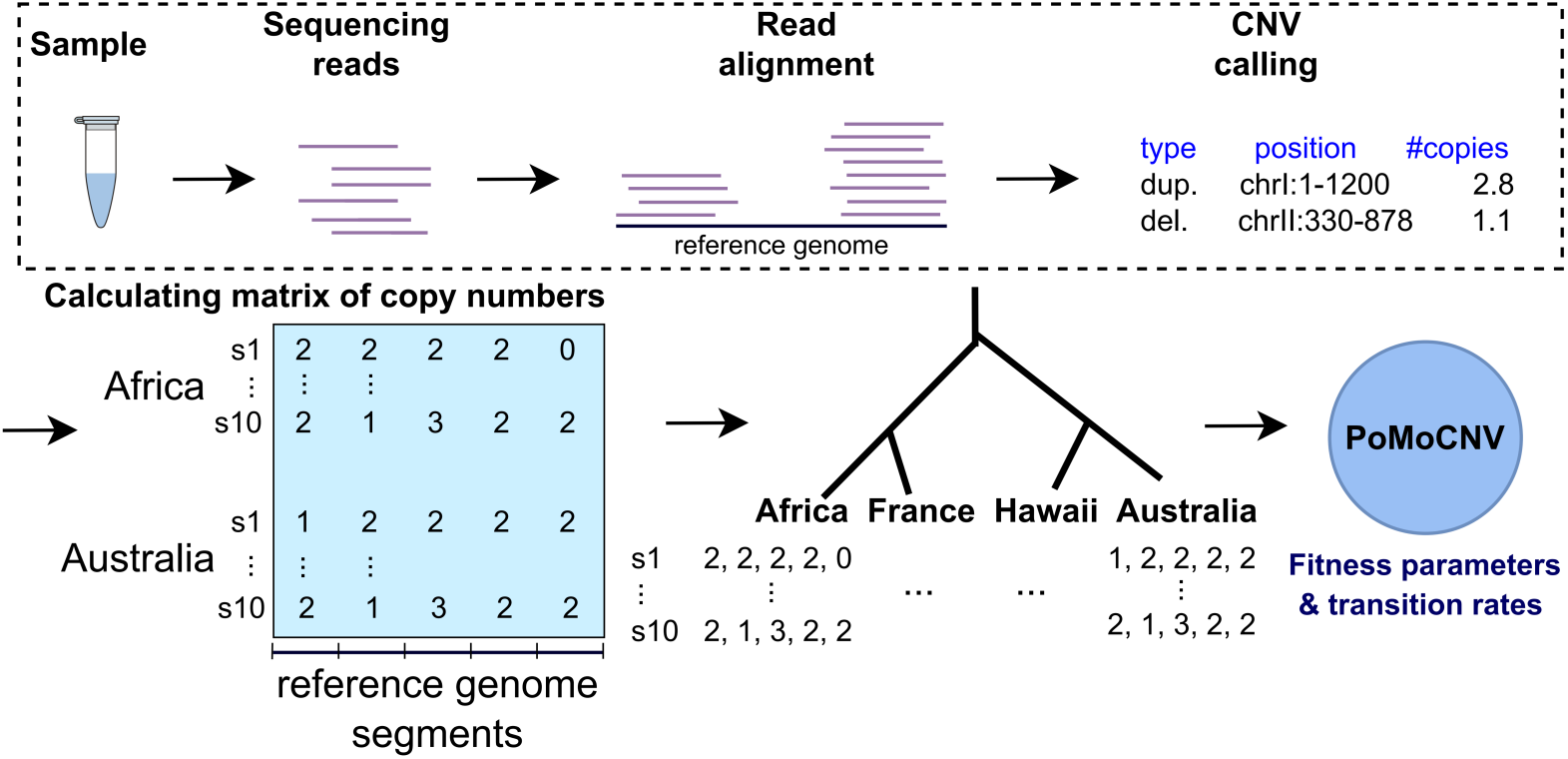
PoMoCNV is employed to estimate fitness parameters and transition rates between different copy numbers in *C. elegans*. Short reads from 10 strains of *C. elegans* for each population (Africa, France, Hawaii, and Australia) are aligned to the N2 reference genome, and used to infer copy numbers for 1kb genomic segments. Utilizing the phylogenetic tree of populations and estimated copy numbers, PoMoCNV was utilized to infer the evolutionary parameters governing CNV evolution along branches.

Our proposed method, PoMoCNV, infers the fitness parameters and transition rates associated with different copy numbers along branches in the phylogenetic tree, tracing back in time. With 4 populations of 10 *C. elegans* strains at the leaves of the tree, the copy number evolution of genomic segments is modeled backward in time to the ancestral population (root node) with a Moran birth-death model. In each generation, one strain is randomly selected to reproduce and transmit its copy number allele to an offspring, while another strain is randomly selected, and its copy number allele is removed from the population. In PoMoCNV, the likelihood of this birth-death process is modeled per genomic segment, taking into account the copy number (allele) fitness and frequencies. For more computational details see Methods section.

We assessed the robustness of the estimated parameters by utilizing 100 bootstrap samples from genomic segments. Figure 2a presents boxplots depicting the estimated transition rates between copy numbers across 100 bootstrap samples. As Figure 2a illustrates, transitions from each copy number to other copies – in close proximity to the original copy – exhibit the highest rates. For instance, transitions from copy number 1 predominantly occur to copy number 0 or 2, from copy number 2 to 1 or 3, and from copy number 3 to 2 or 1. Furthermore, transitions leading to higher copy numbers exhibit higher rates, such as *↵*_12_ *> ↵*_10_, *↵*_23_ *> ↵*_21_, and *↵*_32_ *> ↵*_31_. This preference for copy gains is likely driven by the potential benefits associated with increased gene dosage, while copy losses are generally more detrimental to the organism (Chunduri et al., 2022; Gonzalez et al., 2019; Sung et al., 2016). This suggests that the processes driving copy number changes are influenced by selective pressures favoring the acquisition of additional gene copies over the loss of copies. Figure 2b presents both a violin plot and a box plot illustrating the distribution of the fitness loss per copy deletion (S) across 100 bootstrap samples of genomic segments. The plot reveals that the first quartile (Q1) and third quartile (Q3) of the fitness loss fall within the interval of (0.36, 0.364). The tight clustering of the data points within this interval indicates a relatively low variability in the fitness loss, reinforcing the robustness of the observed trend.

**Figure 2.**
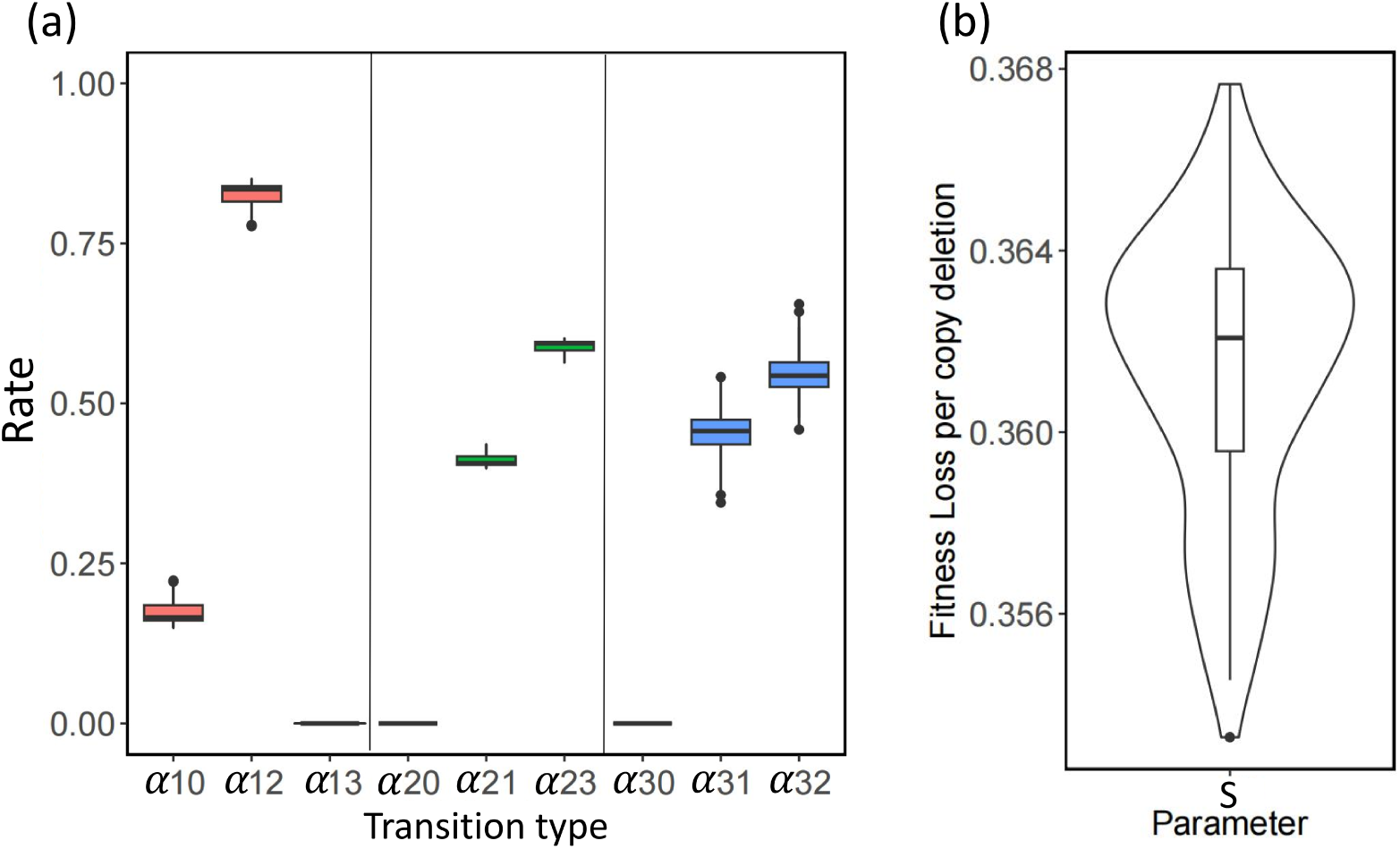
The relative transition rates and fitness loss per copy deletion in genomic segments, in C. elegance. (a) PoMoCNV calculates relative transition rates, which represent the frequency of copy number changes within a genomic segment per generation. (b) PoMoCNV also estimates the fitness loss that occurs with each copy number deletion in a genomic segment.

### CNV evolution rate in open and closed chromatin segments

Understanding the CNV evolution rate in open (accessible) and closed (less accessible) chromatin segments is of significant importance in genomics research. The chromatin state of a gene regulatory segment, whether it is open or closed, plays a crucial role in regulating gene expression (Miyamoto et al., 2018; Wong et al., 2023; Yoshida et al., 2019) and determining the functional consequences of CNVs. Then, investigating the CNV evolution rate specifically in open and closed chromatin segments, (i) furthers our insights into the dynamics of genomic changes and their impact on gene regulation, (ii) sheds light on the interplay between genetic and epigenetic factors (Shi et al., 2020), (iii) elucidates the underlying mechanisms driving CNV formation and selection, providing valuable information for understanding the genetic basis of diseases and the evolutionary processes shaping genomic diversity.

For this purpose, we utilized ATAC-seq data to identify the open and closed chromatin segments (Daugherty et al., 2017; Jänes et al., 2018; Thibodeau et al., 2021). The ATAC-seq peak data from *C. elegans* (Jänes et al., 2018), includes 42,245 peaks with an average width of 152 bp. Collectively, these peaks cover approximately 6.4% of the *C. elegans* genome. In order to investigate the evolution of copy numbers, the genome was divided into segments of 50 bp. Each segment was then annotated based on its overlap with either open or closed chromatin regions derived from ATAC-seq peaks. Following that, we have taken 100 bootstrap samples from the open and closed chromatin segments independently to examine the evolution of CNVs in each segment. Figure shows the boxplots for the transition rates and fitness parameters estimated using PoMoCNV, specifically for the open and closed chromatin segments. The observations presented in Figure 3a indicate that open chromatin segments (depicted in blue) exhibit higher transition rates with increased copy numbers compared to closed chromatin segments (depicted in red). Specifically, the values of *↵*_12_, *↵*_23_, and *↵*_32_ are higher for open segments in comparison to the closed segments. Conversely, open chromatin segments (blue) demonstrate lower transition rates with decreased copy numbers compared to closed chromatin segments (red). In this case, the values of *↵*_10_, *↵*_21_, and *↵*_31_ are lower for open segments relative to the closed segments. Additionally, as depicted in Figure 3b, the fitness loss per copy deletion (S) is notably higher for the open chromatin segments compared to the closed segments. According to the Wilcoxon rank sum test, the difference in the fitness loss per copy deletion between open and closed segments is statistically significant at the alpha=0.05 level.

**Figure 3.**
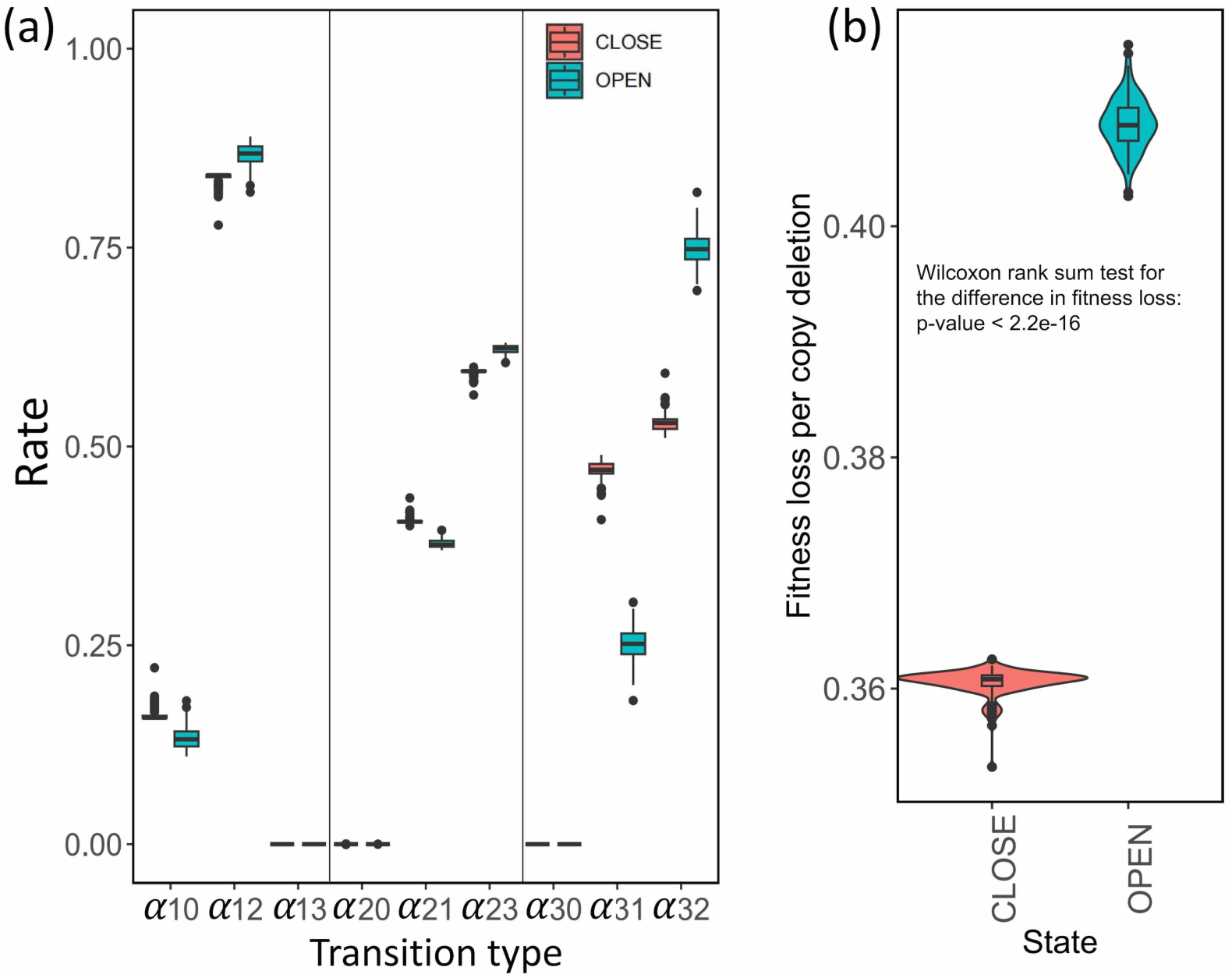
The relative transition rates and fitness loss per copy deletion in open and closed genomic segments in *C. elegans*. PoMoCNV calculates the relative transition rates (a) and estimates the fitness loss resulting from each copy deletion in both open and closed segments (b).

It is important to note that the majority of functional segments, such as enhancers and promoters, primarily reside within the open chromatin segments (Cusanovich et al., 2018; Denny et al., 2016; Ernst and Kellis, 2010, 2012; Klemm et al., 2019; et al., 2012). These functional segments play a crucial role in transcriptional regulation by acting as binding sites for Transcription Factors (TFs). Specifically, TFs facilitate the spatial proximity between enhancers and promoters, enabling the precise regulation of downstream genes in the three-dimensional (3D) genomic landscape (Luo et al., 2022; Pliner et al., 2018). Consequently, any loss of copy numbers within these open chromatin segments can have a more profound impact on the genome. These copy losses in functional segments disrupt the intricate regulatory interactions and can lead to significant impairments in gene expression, potentially resulting in more severe consequences for genomic stability and overall functionality.

### CNV evolution in the synthetic *C. elegans* data

The performance of PoMoCNV is further assessed in CNV datasets from a recent study (Konrad et al., 2018), in which a mutation accumulation (MA) framework was employed to investigate the emergence rate of gene CNVs under varying selection intensities in *C. elegans*. These MA lines were derived from a single hermaphrodite ancestor (N2). In each generation, a bottlenecking process was implemented, resulting in a reduced population size of either *N* = 1, *N* = 10, or *N* = 100 hermaphrodites. Specifically, only one, ten, or one hundred hermaphrodites were selected to reproduce and pass on their genetic information to the next generation, creating the MA lines *N* = 1, *N* = 10, or *N* = 100. After repeating this bottlenecking process for over 400 generations, 18, 40, and 25 descendant individuals from the final generations of each MA line were selected for performing oaCGH (oligonucleotide array comparative genome hybridization) experiments. The aim of these experiments was to assess the presence of copy gains and losses in genomic segments under neutrality (*N* = 1) and with increasing selection intensity (*N* = 10, *N* = 100).

The CNV calls for 18, 40, and 25 *C. elegans* individuals, respectively, from bottleneck sizes *N* = 1, *N* = 10, and *N* = 100, were obtained from a recent study (Konrad et al., 2018). *C. elegans* individuals from each bottleneck size were clustered based on similarities in their copy number patterns using Ward’s method and the heatmap.2 package in R. Supplementary Figure S1 displays the heatmaps showing copy numbers in genomic segments with CNVs and the dendrograms representing similarities among *C. elegans* individuals from each bottleneck size. In Supplementary Figure S1, individuals with similar copy gain or copy loss patterns are closer to each other in the dendrogram. The *C. elegans* individuals from each bottleneck size were further divided into four populations, each consisting of five individuals with similar copy gain or loss patterns based on the Ward’s clustering results. In bottleneck sizes *N* = 10 and *N* = 100, there are no common individuals among four populations. However, in *N* = 1 with 18 individuals, two individuals are shared between two populations to create four populations of size five. To assess the sensitivity of the results to the *C. elegans* individuals included in the constructed populations, we performed 100 bootstrap repeats to sample similar individuals using Ward’s score and assigned them to the same population. For each bottleneck size (*N* = 1, *N* = 10, *N* = 100), we used PoMoCNV to estimate the fitness loss per copy deletion from each bootstrap repeat with four populations of five *C. elegans* individuals. Figure 4 presents boxplots illustrating the fitness loss per copy deletion for *N* = 1, *N* = 10, and *N* = 100. As Figure shows, the estimate of the fitness loss per copy deletion statistically differs among bottleneck sizes *N* = 1, *N* = 10, and *N* = 100, based on the Wilcoxon rank sum test. Moreover, with an increase in the bottleneck size, we observed a lower fitness loss per copy deletion, which allows for an elevated fitness and selection rate for copy gains and losses in genomic segments. Due to the limited number of CNV calls observed in the MA experiments with bottleneck sizes *N* = 1, *N* = 10, and *N* = 100, specifically with 41, 28, and 14 duplication calls, and 39, 16, and 13 deletion calls shared among the sampled *C. elegans* individuals, the estimation of transition rates between different copy numbers is not reported for this dataset.

**Figure 4.**
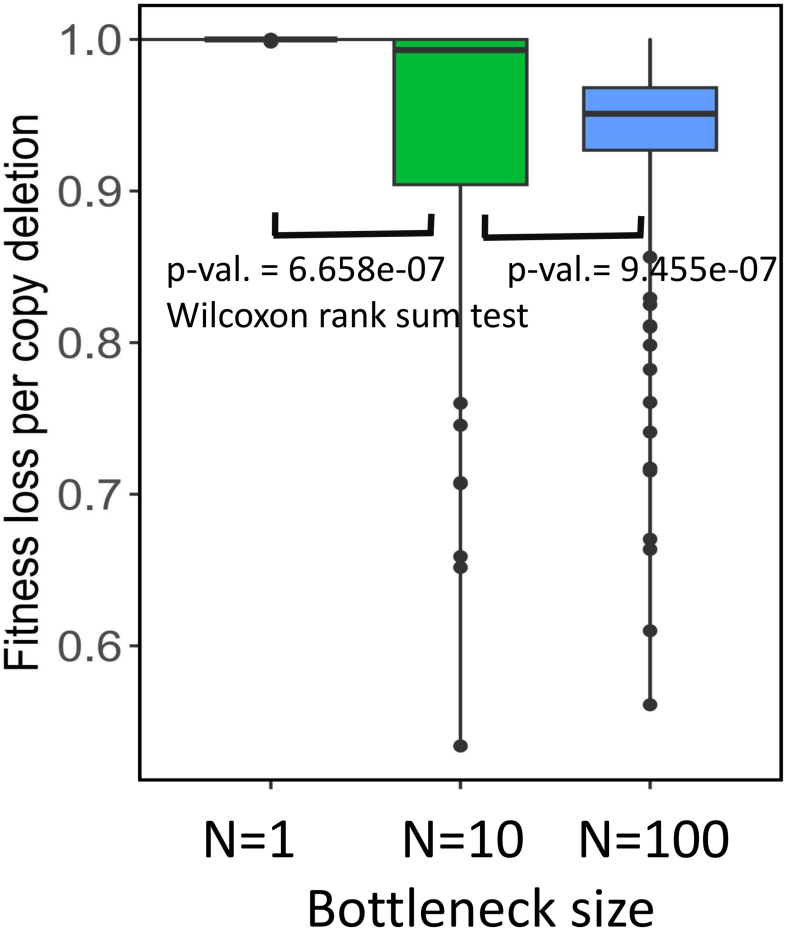
Fitness loss per copy deletion estimated with PoMoCNV for three *C. elegans* datasets with bottleneck sizes *N* = 1, *N* = 10, and *N* = 100. Estimations were performed using 100 bootstrap repeats, dividing *C. elegans* individuals into four populations of size five for each bottleneck size. Individuals within each population share similarities in copy gains and losses compared to the N2 reference genome.

## Discussion

It is generally assumed that the Dynamics of CNVs in populations, mirroring that of SNPs, is shaped by genetic drift and demographic history, as well as selection (Coop et al., 2009; Hollox et al., 2022). In addition, given the large disruptive effects of structural variations on phenotype, CNVs are expected to be non-uniformly distributed across the genome and lay away from functional segments (Conrad and Hurles, 2007). The distribution of CNVs in human genome seemingly abides by these expectations (Sudmant et al., 2015). However, the pattern of CNV distribution does vary from species to species. For example, in an analysis of CNVs in two dairy cattle breeds found a higher overlap between segments of CNVs and genes than expected (Lee et al., 2020).

The fundamental obstacle to a thorough and satisfying evolutionary account of CNVs in a given species is the plethora mechanisms – i.e., retrotransposons, non-allelic homologous recombination, and non-homologous end joining – that can result in CNVs of different size and functions. Our knowledge of demographic history of a lineage, which inevitably leaves its indelible fingerprints on the frequency of CNVs, is generally based on a speculated ancestral state that is the result of comparison with a related lineage (Conrad and Hurles, 2007).

*PoMoCNV* is an attempt to infer aspects of the evolutionary dynamics of CNVs from population genomic data. In the case study of the MA experiment on *C. elegans* (Konrad et al., 2018), given the known ancestral state and varying levels of genetic drift, imposed via varying the bottleneck size in the experiment, enabled us to comprehensively examine the ability of *PoMoCNV* to infer the selective values of CNVs in contrast to our expectation. Based on the available data on chromatin accessibility in *C. elegans*, *PoMoCNV* indicates that CNVs tend to emerge, propagate, and contract more easily in closed chromatin regions. Moreover, the change in copy number in these regions, which are less rich in functional elements, appears to have a comparatively less detrimental effect than CNVs occurring in open chromatin regions. In addition, the fitness effects of CNVs, as inferred by *PoMoCNV*, reflect the interplay between selection and drift, that is, the inferred negative effect of CNVs on fitness increases in concert with the increase in population size, and consequently the increase in the efficacy of natural selection.

## Materials and Methods

In this study, we model the evolution of CNVs along a phylogenetic tree relating populations of individuals. The model considers four copy numbers for a genomic segment – homozygous deletion, heterozygous deletion, normal, and duplication. The evolutionary dynamics are governed by selection coefficients associated with each copy number (allele), along with transition rates between alleles. The transition rate refers to the probability that a genomic segment will transition between two copy numbers during reproduction. For instance, the transition rate may specify the chance that a normal segment becomes duplicated. The transition rates are influenced not only by the frequencies of each copy number in the population, but also by the fitness effects associated with the copy number of each genomic segment. Estimating the underlying fitness parameters and transition rates will shed light on the selective forces and molecular mechanisms generating and propagating CNVs during coursee of evolution.

This integrative approach advances existing literature which has largely focused on either sequence features or population frequencies in isolation. It will help address long-standing questions regarding CNV evolutionary roles, including whether they serve as reservoirs of functional variation or transient neutral changes.

Overall, this research exemplifies how modeling the interplay between selection and copy number changes can further our understanding of genetic variation dynamics and evolution. By leveraging information on population-level copy number (allele) frequencies at the different present-day identities/species, our model aims to disentangle the relative contributions of selection and transition in driving observed CNV patterns.

### PoMoCNV’s framework

We assume that the species are related in a phylogenetic tree and we use the Moran model (De Maio et al., 2013; Nowak, 2006) to estimate the evolution of copy numbers (alleles) along branches in a phylogenetic tree with p=4 populations represented as four leaves of the tree, each including *N*=10 individuals, see Figure 5. For each individual, the model considers four possible strategies (copy numbers) that a gene or genomic segment can adopt – heterozygous and homozygous deletions, normal, and duplication – which are denoting copy numbers 0, 1, 2 and 3, respectively. The population evolves via a stochastic birth-death process. In the birth-death process at each time step, one individual is randomly selected to reproduce and transmit all its copies (allele) to an offspring. Concurrently, another individual is randomly chosen to die and be removed from the population. The reproducing and dying individuals each have their defined copy numbers for all genomic segments. This cycle of reproduction and death, in combination with fitness values, allows the copy numbers to evolve over generations, as individuals with different copy number profiles are randomly chosen to pass on their genetic material or die out each time step.

**Figure 5.**
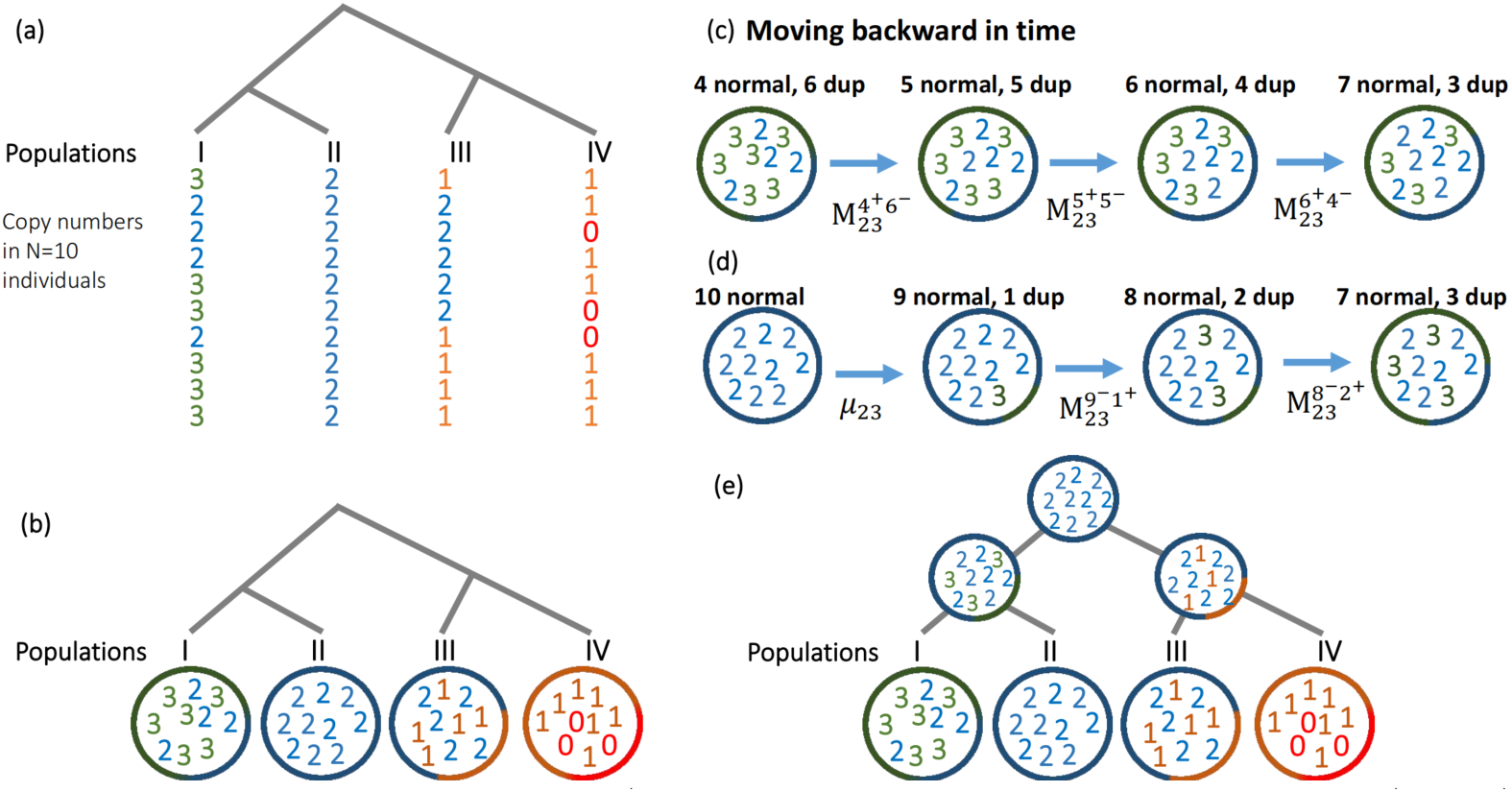
Pipeline for PoMoCNV (POlymorphism-aware phylogenetic MOdel (PoMo) for CNV). (a) The copy numbers (alleles) of 10 individuals from four *C. elegans* populations are displayed along the phylogenetic tree for a specific genomic segment. (b) Each genomic segment within each *C. elegans* population is associated with a PoMoCNV state based on the copy number distribution. (c) In multiallelic states, which include distinct copy numbers within a population, PoMoCNV utilizes a Moran model with a state transition rate matrix (M) to simulate backward movement between different states over time. (d) In monoallelic states, where only one copy number is observed in a population, PoMoCNV utilizes a transition rate (*↵*) to introduce a new copy number (allele) for a potential biallelic population at the inner nodes of the phylogenetic tree. (e) The Felsenstein pruning algorithm is utilized by PoMoCNV to aggregate the likelihoods across all potential states at the inner nodes. This allows for the estimation of the most probable fitness and transition rates associated with copy numbers.

### States and transition probabilities

The model considers populations of *N* diploid individuals, and each individual is with a copy number (allele) for each genomic segment. At each genomic segment, up to two alleles are allowed in the population in our modeling. The possible alleles are denoted *I* and *J*, representing the four copy numbers 0, 1, 2 and 3. The allele frequency distribution can be summarized by the notation *{i, j}*, where *i* and *j* refer to the counts of alleles *I* and *J*, respectively, in the population of *N* individuals. This allows representation of all possible allele configurations ranging from monoallelic states with all *I* (*{*0*,N}*) or *J* (*{N,* 0*}*), along with intermediate biallelic states like *{*1*,N -*1*}*, *{*2*,N -*2*}*, . . ., *{N -*1, 1*}*. For a population with *N* = 10 diploid individuals and four possible allele types, there are a total of 4 + 6 *⇥* (*N -* 1) = 58 distinct allele frequency states, encompassing the spectrum from monomorphic to polymorphic. By tracking transitions between frequency states over generations, we can infer evolutionary dynamics and estimate key parameters that govern the transition and selection processes affecting copy number variation. This analysis specifically focuses on copy number variation at the genomic segment within the branches of the phylogenetic tree that connects different populations.

For a biallelic genomic segment with copy numbers *I* and *J* existing in the population, the Moran model allows calculation of transition probabilities between allele frequency states. Specifically, the probability of transitioning from state *{i, j}* to *{i* + 1*,j -* 1*}* in one generation can be denoted as *M ^i^*^+^*^,j-^* and derived as:

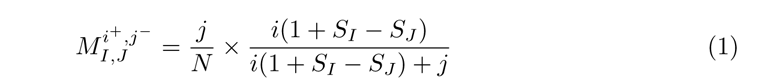

In this formulation, *^j^* represents the probability of randomly selecting an individual with allele *J* to die, while the term *^i^*^(1+^*^SI -SJ^*^)^ *I J* gives the probability of choosing an individual with allele I to reproduce, proportional to the fitness advantage S_I_ of allele I over J. Similarly, the reverse transition probability *M* ^*i-,j+*^ from state *{i, j} to {i - 1,j + 1}* is:

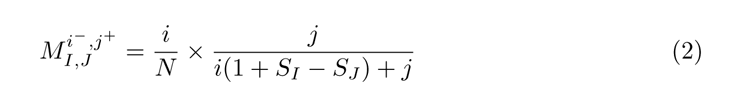

These transition probabilities depend on both the current allele frequencies i and j, as well as the selection coefficients *S*_I_ and *S*_J_ associated with each allele. This enables modeling of how selection and genetic drift shape CNV evolutionary dynamics.

The transition from a monoallelic state to a biallelic state can happen when a new copy number appears in the population which could be the result of a ”transition” event, i.e. change in copy number, governed by a ”mutation” rate. Specifically, the probability of transitioning from a state with N individuals having a copy number *I* to another state characterized by copy numbers *I* and *J*, where the frequencies are {*N* - 1, 1}, is calculated as follows:

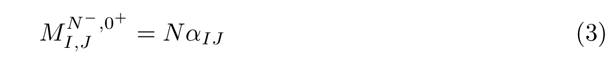

where ↵_IJ_ represents the transition rate from an individual with copy number *I* to copy number *J* within a single generation. The relative relationships among fitness parameters of copy numbers are considered as *S*_0_ = 1 - 2 *S*, *S*_1_ = 1 - *S*, *S*_2_ = 1, and *S*_3_ = 1 - 0.5*S*.

In this formulation, we designate genomic segments with normal copies as a reference, assigning them a fitness value of *S*_2_ = 1. The parameter S, which can range within the interval (-0.5, 0.5), represents the overall decrease or increase in fitness for genomic segments with each copy loss. To estimate this parameter, along with other model parameters, the data likelihood is maximized with Felsenstein pruning algorithm, as implemented in the HyPhy package (Pond et al., 2004). The fitness change in genomic segments with amplified copies (S_3_) is relatively lower compared to copy losses, as it is not as detrimental to the functionality of the genomic segments (Chunduri et al., 2022; Gonzalez et al., 2019; Sung et al., 2016). For example, spinal muscular atrophy (SMA) is caused by homozygous deletion of SMN1 (Lefebvre et al., 1995), and individuals with the mildest form of SMA often possess three or more copies of SMN2 (Feldkötter et al., 2002; Mailman et al., 2002), enabling their survival into adulthood. Moreover, homozygous deletions exhibit a higher occurrence within genomic regions characterized by a lower gene density, relative to other types of genetic variations (Girirajan et al., 2011). In other complex diseases such as autism, schizophrenia, bipolar disorder, amyotrophic lateral sclerosis, attention deficit hyperactivity disorder (ADHD), and Tourette syndrome large deletions play a crucial role (Girirajan et al., 2011). In individuals experiencing developmental delay, the occurrence of deletions is more prevalent compared to reciprocal duplications (Girirajan et al., 2011). However, despite this, copy gains still impose a burden on the genome by increasing its size. Consequently, genomic segments with copy gains will exhibit lower fitness compared to segments with normal copies.

## Supplementary Material

Supplementary files are uploaded separately.

## Data Availability

The HyPhy codes and data analyzed in the article are available on the GitHub page of the project ”PoMoCNV” at the following URL: https://github.com/CompBioIPM/PoMoCNV.

## Supplementary file

**Supplementary Figure S1:**
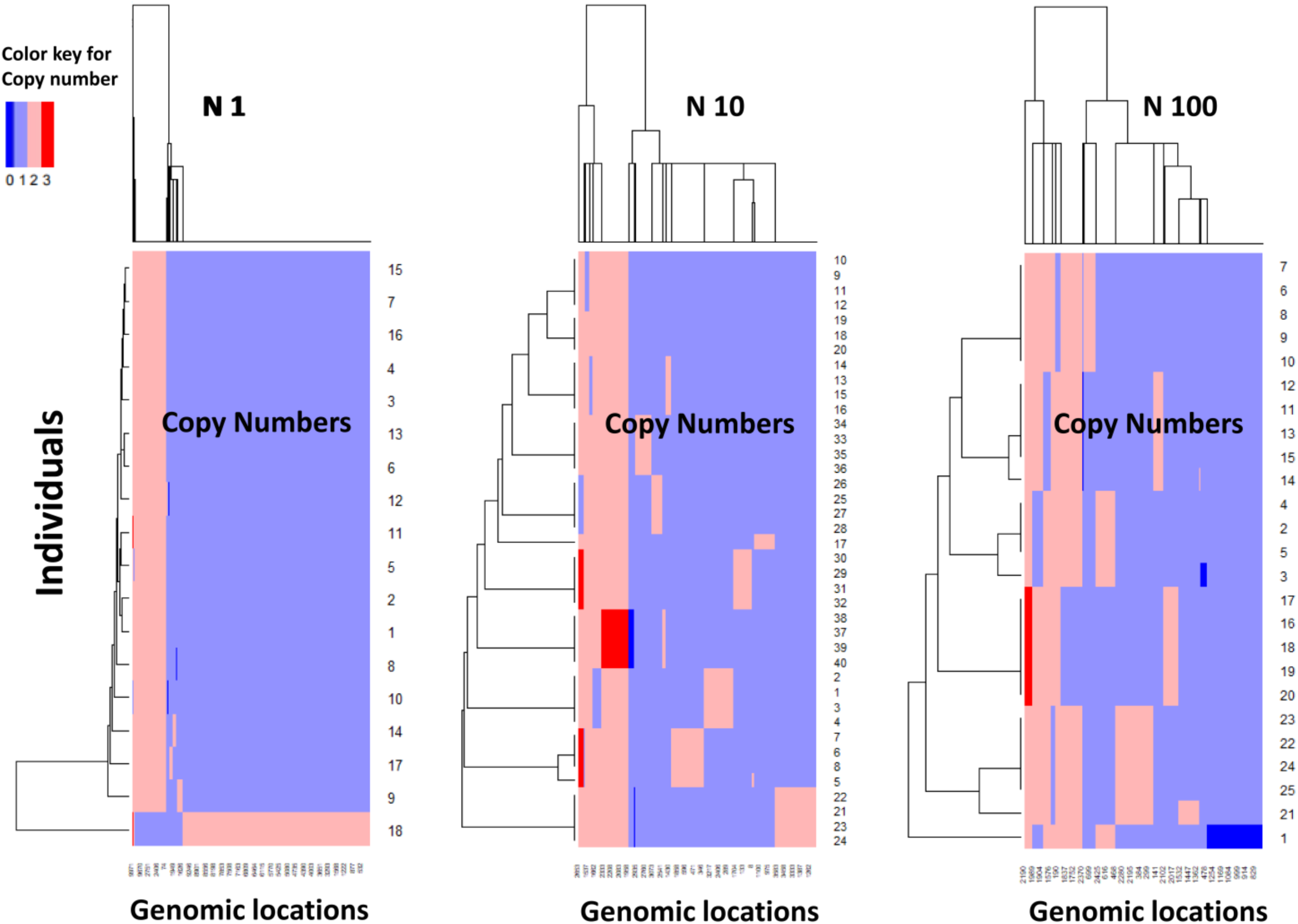
CNV profile of *C. elegans* individuals in mutation accumulation (MA) experiments. The figure displays the genomic locations with CNVs, along with their corresponding copy numbers represented in distinct colors, for 20, 40, and 25 samples of *C. elegans* individuals in the N=1, N=10, and N=100 experiments, respectively.

